# Associations between parenting and cognitive and language abilities at age 2 depend on prenatal exposure to disadvantage

**DOI:** 10.1101/2024.03.25.586610

**Authors:** Shelby D. Leverett, Rebecca G. Brady, Ursula A. Tooley, Rachel E. Lean, Rebecca Tillman, Jillian Wilson, Michayla Ruscitti, Regina L. Triplett, Dimitrios Alexopoulos, Emily D. Gerstein, Tara A. Smyser, Barbara Warner, Joan L. Luby, Christopher D. Smyser, Cynthia E. Rogers, Deanna M. Barch

**Author notes:** **Corresponding Author:** Shelby Leverett 4525 Scott Avenue, St Louis MO 63310. **Data Sharing Statement**: Data is available through the National Data Archive. Study data was presented as a poster at FLUX Conference 2023.

## Abstract

**Objective:** To investigate whether parenting and/or neonatal brain volumes mediate the associations between prenatal social disadvantage (PSD) and cognitive/language abilities; and whether these mechanisms vary by level of disadvantage.

**Study Design:** Pregnant women were recruited from obstetric clinics in St Louis, Missouri. PSD encompassed access to social (e.g., education) and material (e.g., income-to-needs, health insurance, area deprivation, and nutrition) resources during pregnancy. Neonates underwent brain magnetic resonance imaging. Mother-child dyads (N=202) returned at age 1 for parenting measures and at age 2 for cognition/language assessments (Bayley-III). Generalized additive and mediation models tested hypotheses.

**Results:** Greater PSD was nonlinearly associated with poorer cognitive/language scores. The relation between parenting and cognition/language was moderated by PSD, such that supportive and non-supportive parenting behaviors only related to cognition/language in children with low PSD. Further, parenting mediations differed by level of PSD, such that both supportive and non-supportive parenting mediated PSD-cognition/language associations in children with low PSD, but not in children with high PSD. PSD-associated reductions in neonatal subcortical grey matter (β=.19, *q*=.03), white matter (β=.23, *q*=.02), and total brain volume (β=.18, *q*=.03) were associated with lower cognition, but they did not mediate PSD-cognition associations.

**Conclusions:** Parenting moderates and mediates associations between PSD and early cognitive and language development, but only in families with lower levels of social disadvantage. These findings, while correlational, suggest that there may be a critical threshold of disadvantage, below which mediating or moderating factors become less effective, highlighting the importance of reducing disadvantage as primary prevention.

## INTRODUCTION

Social disadvantage (SD), or lack of access to material and social resources, during childhood is consistently associated with lower cognition and language (CaL) abilities.^1–3^ This disparity strongly contributes to a socioeconomic achievement gap^4^ that begins in childhood and is sustained or even increases with age.^5^ There is evidence that socioeconomic-related variability in CaL abilities emerges early in development,^6,7^ with most recent evidence linking variability in *prenatal* SD (PSD) to language.^8^ Before successfully identifying resilience factors and effective intervention targets, it is critical to better understand mechanisms through which PSD is associated with challenges to early CaL development.

There may be multiple mechanisms through which PSD influences CaL. One may include the impact of PSD on prenatal fetal brain development, evident in structural or functional alterations at birth. Greater PSD has been associated with altered brain structure and function in neonates.^9–11^ Moreover, in older children, such alterations both have been correlated with developmental abilities like CaL^12–15^ and may mediate the relation of *postnatal* SD to CaL.^15–17^ Thus, these neural phenotypes *at birth* may also mediate the relationship between *prenatal* SD and CaL. Another possible mechanism may be parenting behaviors-which are influenced by correlates of SD^18–20^ and are known to influence offspring academic outcomes.^21–25^ Moreover, parenting has been shown to mediate the relations of postnatal SD to CaL abilities in older children^26–28^ but has yet to be examined as a similar mediator of *prenatal* SD and CaL. However parenting is particularly influential in early life,^29,30^ and constitutes a potentially malleable intervention target, thus is an important mechanism to examine.

Finally, a complete mechanistic understanding requires mechanistic specificity. It is possible that mechanisms of PSD-CaL associations vary by levels of disadvantage. In fact, several pathways shown to mediate disadvantage-outcome associations demonstrate differential relations in high versus low disadvantage conditions.^31,32^ This mechanistic specificity is a critical phenomenon to consider, as it increases explanatory power^31^ and creates insights for tailored preventative interventions to be maximally effective.

In this prospective, longitudinal study, we sought to better understand mediating and moderating pathways between PSD and offspring CaL outcomes at age two years. We first sought to replicate the known association of SD-CaL in our study sample. Then we tested the following novel hypotheses: associations between PSD and CaL would be mediated by (1) PSD-associated variation in brain volumes at birth and (2) parenting behaviors; and (3) disadvantage moderates the relationships between identified mediators and outcomes (e.g., relationships between parenting and cognition/language vary across levels of disadvantage).

## METHODS

### Participants and Procedures

Data were collected as part of the Early Life Adversity Biological Embedding (eLABE) study.^33,34^ The study enrolled 395 healthy pregnant women and their singleton offspring (see Supplement for enrollment criteria). Women were over-sampled from clinics serving low-income populations to enrich the sample for exposure to poverty. Women were enrolled early in pregnancy and completed assessments in each trimester. Neonates underwent brain MRI scans.

Mother-child dyads returned for follow-up assessments at ages 1 and 2 years. Written informed consent was obtained from mothers after describing study protocols and prior to participation. All protocols were approved by the Institutional Review Board at Washington University in St. Louis.

### Measures

#### Prenatal Social Disadvantage

As described in Luby et al,^33^ PSD is a previously validated latent construct that includes variables relating to material and social resources during pregnancy: income-to-needs ratio, Area Deprivation Index, health insurance status, maternal education, and maternal nutrition. Higher PSD scores represent more disadvantage. See *Supplement* for more information.

#### Bayley-III

The Bayley Scales of Infant and Toddler Development, Third Edition (Bayley-III) is a gold-standard assessment of cognitive, language, and motor development for children aged 1-42 months with good reliability and validity.^35^ The Bayley-III was administered by highly trained psychometricians. The primary dependent measures were composite scores for cognition, language, and motor abilities at year 2 follow-up, wherein higher scores indicate better performance. To explore specificity to CaL associations, motor abilities — which have previously been unrelated to PSD in prior work^36^— were included as a control outcome.

#### Parenting Behaviors

An adapted version of the Parent-Child Interaction Rating Scales (PCIRS)^37^ was used to code parent-child interactions across three videotaped tasks. Parenting behaviors were scored by trained raters on a 7-point scale and averaged across tasks. A Supportive Parenting Behaviors composite (α=.861) averaged scores for sensitivity and positive regard. A Non-Supportive Parenting Behaviors composite (α=.726) averaged scores for intrusiveness, detachment, and negative regard.

Higher scores represent greater intensity and frequency of the observed behavior. Analyses used parenting behaviors at year 1 to enable mediation analyses with temporal precedence of parenting (year 1) relative to CaL (year 2). See Supplement for more details and reliabilities.

#### Test of Premorbid Function (TOPF)

The TOPF^38^ was used to assess maternal cognitive ability. It is a standardized task in which participants pronounce up to 70 words with irregular grapheme to phoneme translations. Age-normed TOPF standard scores are reported.

#### Magnetic Resonance Imaging (MRI)

MRI data collection and processing were conducted as previously described.^10^ Briefly, all infants were scanned during natural sleep without sedation in the first weeks of life (m=3.2 weeks, sd=1.9, range=0-10) on a Siemens Prisma 3T scanner with a 64-channel head coil to acquire T2-weighted images (TR=3200/4500ms, TE=563ms, 0.8mm^3^ isometric voxels) and spin-echo field maps (TR=8000ms, TE=66ms, 2mm^3^ isotropic voxels, multiband factor=1). Preprocessing pipelines are further detailed in the Supplement. Prior work in this sample^10^ found associations between greater PSD and volumetric reductions in six neonatal brain metrics: total brain, cortical GM, subcortical GM, total WM, and cerebellar volumes and gyrification index (GI).^10^ These six brain metrics were the *a priori* measures of interest.

### Data analysis

#### Covariates

Gestational age (GA), infant sex at birth, and age at follow-up were related to Bayley-III scores and were therefore included as covariates in all analyses. See *Supplement* for statistics.

#### Modeling PSD and child outcomes

There is increasing emerging evidence of nonlinear associations between socioeconomic status and brain development.^16,39,40^ Thus, predictors and covariates were added to generalized additive models (GAMs), a type of nonparametric regression wherein the relationship between the predictor and outcome is not assumed to be linear; allowing either nonlinear or linear relations can be detected and modeled in a data-driven fashion, GAMs return an expected degree of freedom (EDF) which specifies the shape of the “smooth”. The restricted likelihood ratio test (RLRT) (*exactRLRT* function from *RLRsim* package^41,42^ in R) determined whether the nonlinear fit of each smooth was significantly better than the linear fit. If the better fit was nonlinear (RLRT *p*-value <0.05), GAMs were used in all subsequent analyses. To visualize periods of significant change in smooths (e.g., where the dependent variable significantly changes as a function of the independent variable), the 95% confidence interval of the derivative of each significant smooth was calculated and mapped back onto the smooth using the *gratia* package in R.^42–44^ If the nonlinear model was not a better fit (RLRT *p*-value>0.05), linear regressions were used (*stats* package^45^). Effect sizes for nonlinear models were calculated using change in adjusted R^2^-values (ΔAdj. R^2^; the difference between the adjusted R^2^-values of a model with and without the predictor(s) of interest), whereby positive values indicate that the predictor improves the model fit.

If an outcome was associated with PSD, we tested whether that association was significant over and above the influence of maternal cognitive ability by adding TOPF to the model.

#### Neural Mediations

Regressions with covariates of interest tested whether each of the six *a priori* brain metrics related to primary outcomes. When birth brain metrics predicted 2-year outcomes, we tested whether each metric mediated the relations between PSD and associated outcomes.

#### Parenting Behavior Mediation

Regressions with covariates tested whether: (1) PSD related to parenting behaviors, (2) parenting behaviors related to outcomes, and (3) parenting behaviors mediated and/or moderated the relationships between PSD and outcomes. Analyses were conducted separately for supportive and non-supportive parenting behaviors.

#### Disadvantage Moderation

Moderation was tested by adding an interaction between PSD and parenting composites to each outcome. Nonlinear moderations preclude follow-up with simple slopes. Therefore, significant interactions were probed by examining associations (with GAMs) between one independent variable and the outcome separately in high vs low disadvantage groups, as defined by a median split. Finally, for significant mediators of PSD-CaL associations, we tested whether the mediation was significant in high vs low disadvantage conditions, as defined by a median split.

All mediations were conducted with GAM (nonlinear) or ‘lm’ (linear) objects in R (*mediation* package v4.5.0). All analyses used complete cases (listwise deletion). Results for all primary analyses were corrected using the False Discovery Rate corrections calculator.^46^ For correction, tests were grouped by the research question and by the outcome variable. See Supplement for further details.

## RESULTS

Children missing Bayley-III scores were excluded from analyses, resulting in a sample of 202 children. Sample characteristics are detailed in Table 1. Included and excluded children did not differ on any variables in analyses.

**Table 1:** Sample Characteristics.

| **Maternal Characteristics, mean(sd) [range]** |  |
| --- | --- |
| Prenatal social disadvantage latent factor score, *N*=202 | -.06 (0.97) [-2.22-1.55] |
| Income-to-needs ratio | 2.94 (3.1) [0.32-12.15] |
| National Area Deprivation Index | 69.78 (24.05) [7-100] |
| Healthy Eating Index | 58.47 (10.47) [31.70-80.67] |
| Highest level of education, *n* (%) |  |
| *less than 12th grade* | 7 (3.46%) |
| *high school degree/GED* | 44 (21.8%) |
| *some college/vocational school* | 70 (34.6%) |
| *college degree (4 years)* | 25 (12.4%) |
| *graduate degree* | 56 (27.7%) |
| Health insurance status, *n* (%) |  |
| *Medicaid or Medicare* | 69 (34.1%) |
| *Individual or Group Health Insurance* | 113 (60.0%) |
| *VA/Military* | 1 (0.5%) |
| *Uninsured* | 19 (9.4%) |
| Test of Premorbid Functioning, *N*=152 | 94.22 (16.24) [68-124] |
| Supportive Parenting Behaviors at Year 1, *N*=135 | 4.76 (0.89) [2.33-6.33] |
| *sensitivity* | 4.81 (0.95) [2.33-6.67] |
| *positive regard* | 4.71 (0.97) [2.00-7.00] |
| Non Supportive Parenting Behaviors at Year 1, *N*=135 | 1.64 (0.50) [1.00-3.67] |
| *negative regard* | 1.33 (0.52) [1.00-3.67] |
| *intrusiveness* | 2.16 (0.89) [1.00-5.00] |
| *detachment* | 1.41 (0.67) [1.00-4.33] |
| **Child Characteristics (*N*=202), mean(sd) [range]** |  |
| Gestational Age at Birth (weeks) | 37.89 (1.9) [28-41] |
| Age at Year 2 (years) | 2.10 (.14) [1.91-2.64] |
| Sex assigned at birth, *n* (%) | - |
| *male* | 110 (54.5%) |
| *female* | 92 (45.5%) |
| Child race, *n* (%) | - |
| *African American* | 120 (59.4%) |
| *White* | 77 (38.1%) |
| *Chinese* | 1 (0.5%) |
| *Korean* | 1 (0.5%) |
| *Pacific Islander* | 1 (0.5%) |
| *Multiracial* | 1 (0.5%) |
| *other* | 1 (0.5%) |
| Bayley-III Cognition scores at Year 2 | 88.61(12.75) [55-125] |
| Bayley-III Language scores at Year 2 | 89.04(16.69) [59-138] |
| *expressive language* | 8.74 (2.80) [2-19] |
| *receptive language* | 7.47 (3.22). [1-16] |
| Scan Age (weeks) N=186 | 3.2 (1.9)[0-10] |
| Neonatal Brain Volumes (mm<sup>3</sup>), N=186 |  |
| *total cortical GM* | 120303.1 (15939.3) [7318.5 - 161621.8] |
| *total cerebral WM* | 185282.3 (19424.6) [136548.9- 244443.3] |
| *total cerebellum* | 28273.7 (4507.9) [15645.7 - 45875.2] |
| *subcortical grey* | 27115.2 (2703) [20404.8 - 35261] |
| *total brain volume* | 359364.2 (39404.7) [263109.5 - 472884.4] |
| *mean GI* | 1.97 (.10) [1.63 - 2.20] |

### PSD and Primary Outcomes

PSD was non-linearly related to cognition (ΔAdj. R^2^=12.47%, EDF=2.62, RLRT *p=*.02, q<.0001) and language (ΔAdj.R^2^=17.21%, EDF=2.70, RLRT *p<*.0001, *q*<.0001) As shown in Figure 1, greater PSD was associated with poorer CaL scores up to a certain level of PSD, beyond which additional disadvantage was not associated with further decreases in scores. PSD continued to predict cognition (β = -.26, *p=*.037) and language (β = -.33, *p=*.007) even after accounting for maternal cognitive ability. PSD was not related to motor abilities (β = -.064, *q*=.35).

**Fig 1:**
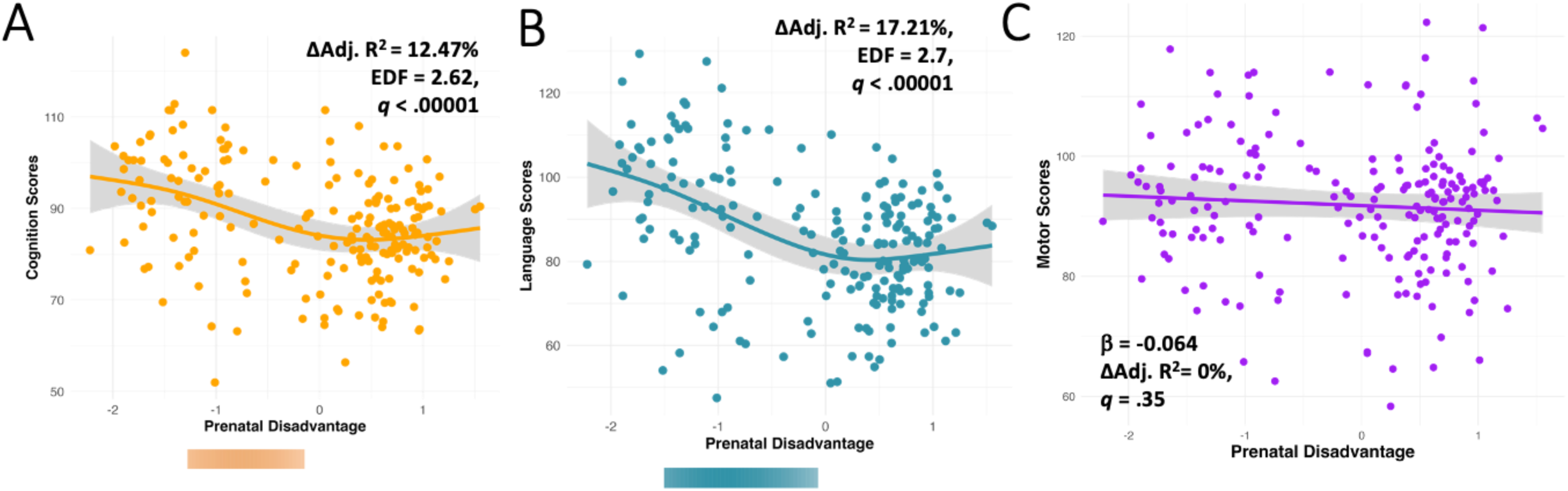
Scatterplots of the relation of prenatal disadvantage to cognition, language, and motor outcomes at Year 2. **Associations between PSD and (A) cognition, (B) language, and (C) motor outcomes at Year 2**. After controlling for sex, gestational age at birth, and age at assessment, greater PSD was nonlinearly associated with poorer cognition and language scores up to a point, beyond which additional PSD was not associated with further decreases in scores. Colored bars below scatterplots represent regions of significant change, where confidence interval of derivative excludes 0, indicating the region where cognition and language are significantly changing in relation to PSD. After controlling for sex, gestational age at birth, and age at assessment, PSD was unrelated to motor scores at year 2. *EDF = expected degrees of freedom*. Δ Adj R^2^ = the difference between the adjusted R^2^-values of a model with and without the predictor of interest, where positive values indicate that the predictor of interest improves the model fit.

### Neural Mechanisms

Reduced subcortical GM, WM, and total brain volumes at birth were linearly (all RLRT *p*-values>.25) associated with lower cognitive scores (eTable 1). No brain metric at birth related to language scores (eTable 1). None of the related brain metrics mediated associations between PSD and cognition or language (eTable 2).

### Parenting Mechanisms

Greater PSD was linearly (RLRT *p’s*=1) associated with fewer supportive parenting behaviors (β = -.64, *q*<.0001) and more non-supportive parenting behaviors (β =.55, *q*<.0001) at year 1 (eFigure 1). Higher levels of supportive parenting behaviors were nonlinearly associated with higher cognition (ΔAdj. R^2^=14.87%, EDF=2.34, RLRT *p=*.03) and language scores (ΔAdj. R^2^=19.01%, EDF=2.35, RLRT *p=*.02), such that the relationship of supportive parenting to outcomes was stronger among higher supportive parenting scores (eFigure 1). Higher levels of non-supportive parenting behaviors were nonlinearly associated with poorer cognition (ΔAdj. R^2^=11.37 %, EDF=4.17, *p=*.027) and language scores (ΔAdj. R^2^=12.31%, EDF=3.03, *p=*.004), such that the relation of non-supportive parenting to cognition and language was strongest at lower levels of non-supportive parenting (eFigure 1).

Further, lower supportive parenting mediated the relation of greater PSD to lower CaL scores (Fig 2), though non-supportive parenting did not (eTable 2).

**Fig 2:**
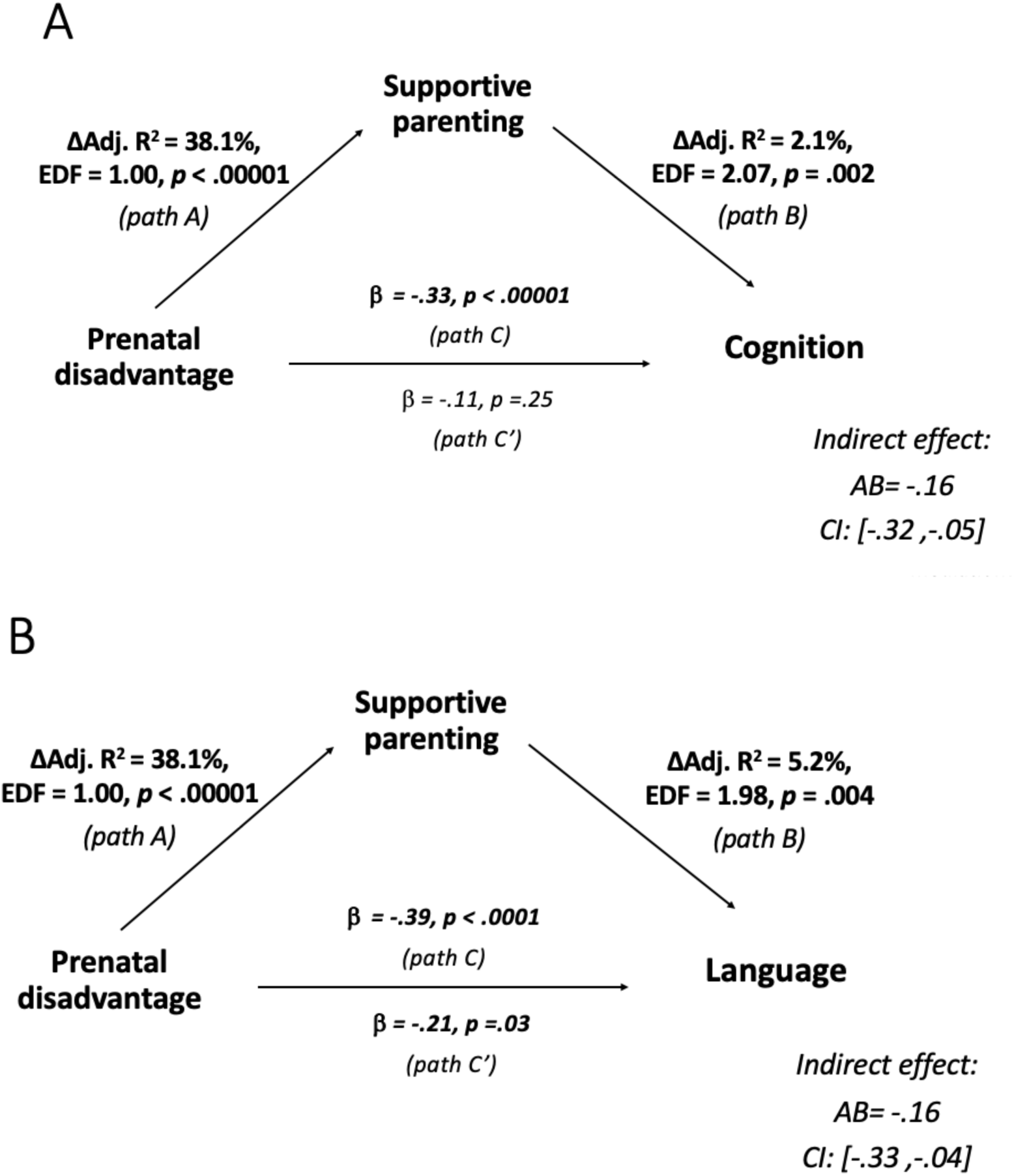
Supportive Parenting Behaviors mediate the relations of prenatal disadvantage to (A) cognition and (B) language. **Mediation models. Standardized effect sizes, p-values and/or confidence intervals for total, direct, and indirect effects. Change in Adjusted R^2^ values (Δ Adj. R^2^), expected degrees of freedom (EDF) and GAM p values for paths A and B, which were modeled with GAMS (RLRT p < .05)**. Path A describes the effect of the independent variable (prenatal disadvantage) to the mediator variable. Path B describes the unique effect of the mediator variable on the dependent variable. Path C’ (direct effect) indicates the unique effect of the independent variable (prenatal disadvantage) on the dependent variable, while controlling for the mediating variable. The indirect effect is the mediation effect. Path C indicates the total effect. Δ Adj R^2^ = the difference between the adjusted R^2^-values of a model with and without the predictor of interest, where positive values indicate that the predictor of interest improves the model fit.

### Disadvantage Moderation

Disadvantage moderated relations of parenting to cognition (supportive interaction: ΔAdj. R^2^=4.2%, EDF=3.00, *q*<.0001; non-supportive interaction: ΔAdj. R^2^=4.4%, EDF=5.54, q<.0001) and to language (supportive interaction: ΔAdj. R^2^=1.6%, EDF=3.00, *q*<.0001; non-supportive interaction: ΔAdj. R^2^=1.9%, EDF=3.18, *q*<.0001) scores (Fig 3). Specifically, follow-up analyses revealed that for children with *lower* levels of PSD, supportive and non-supportive parenting behaviors were respectively positively and negatively associated with cognition (supportive: p=.0002; non-supportive: p=.0049) and language (supportive: p=.00007; non-supportive: p=.0074). However, at higher levels of PSD, the association between parenting and cognition/language was no longer significant (all p’s>.05).

**Fig 3:**
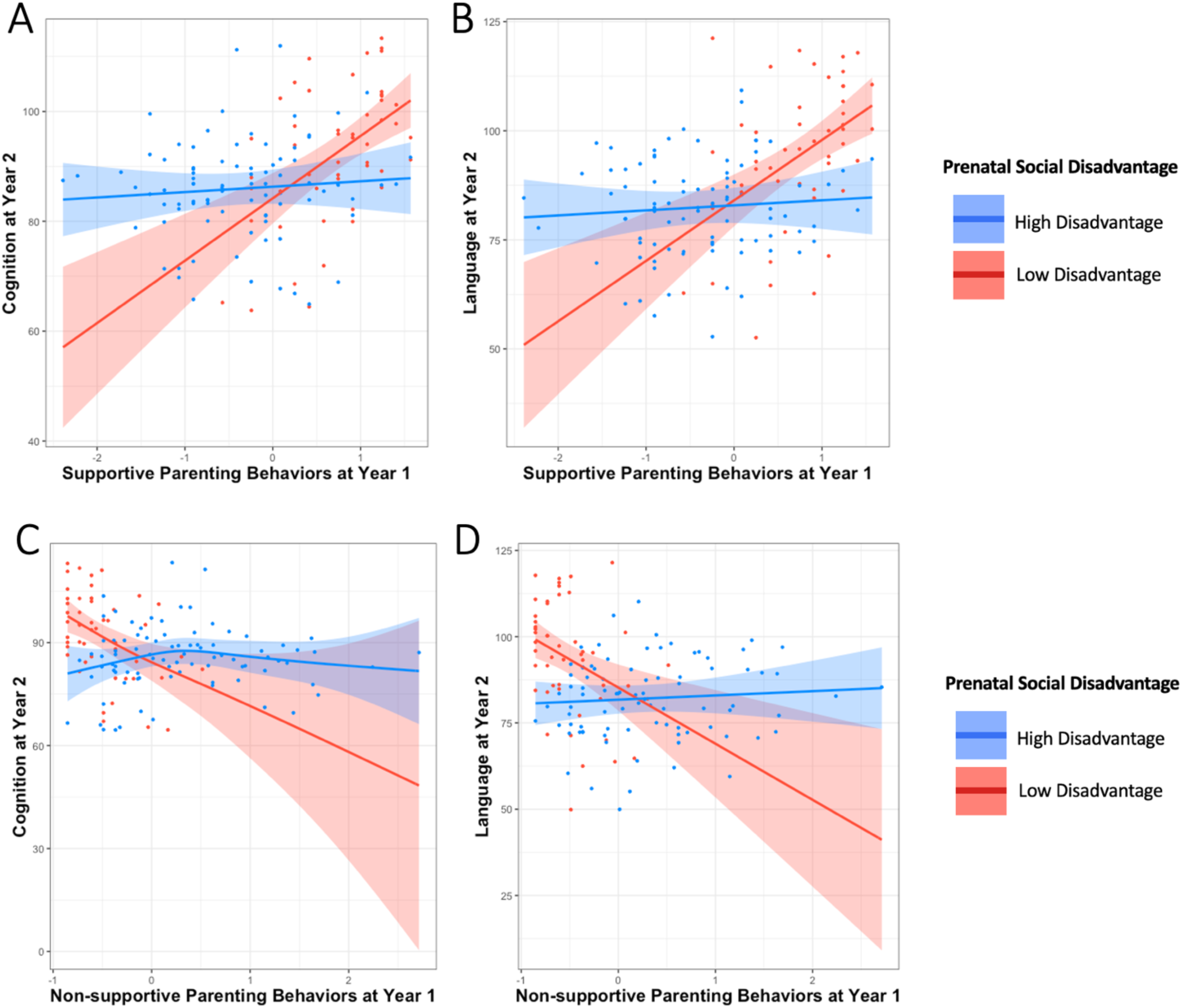
Prenatal Disadvantage Moderates the Association between Parenting Behaviors and Cognition and Language Scores. Interaction of PSD with Supportive Parenting Behaviors to predict year 2 (A) cognition (q<.000001) and (B) language (q<.000001). Interaction of PSD with non-supportive parenting behaviors to predict year 2 (C) cognition (q<.00001) and (D) language (q<.0001). PSD is split at median for visualization, however, all analyses used continuous PSD. Model: *gam(outcome ∼ te(PSD, parenting) + GA + child_sex + age, method = “REML”, fx = T)*

Given the parenting moderation and mediation identified above, exploratory mediation analyses of parenting behaviors were examined separately in high and low levels of disadvantage, defined by a median split of high (N=71) and low (N=64) disadvantage. Parenting (both supportive and non-supportive) mediated the relationship of PSD to cognitive and language outcomes in the low disadvantage group, but not in the high disadvantage group (Table 2).

**Table 2:** Parenting mediations in high disadvantage vs low disadvantage. **Parental behaviors mediate the relations of PSD to cognition and language in the low disadvantaged sample, but not in the high disadvantaged sample. *Standardized effect estimates (μ) for path A, path B, direct and indirect effects.*** *Path A describes the effect of the independent variable (prenatal disadvantage) on the mediator variable. Path B describes the unique effect of the mediator variable on the dependent variable (outcome). Direct effect indicates the unique effect of independent variable (prenatal disadvantage) on the dependent variable, while controlling for the mediating variable. The indirect effect is the mediation effect. Covariates in the model are sex, age at assessment, and gestational age at birth*. ***Bolded rows indicate a significant mediation effect. *p < .05, **p < .01, ***p < .001, ****p < .0001***

| | Mediating Variable | Outcome | path A<br>$\beta$ | path B<br>$\beta$ | Direct effect<br>$\beta$ | indirect effect | 95% CI Indirect Effect | |
| --- | --- | --- | --- | --- | --- | --- | --- | --- |
|  |  |  |  |  |  |  | Lower | Upper |
| High Disadvantage (N=71) | supportive parenting behaviors | cognition | -0.25 | 0.09 | 0.21 | -0.023 | -0.17 | 0.07 |
|  | supportive parenting behaviors | language | -0.25 | 0.12 | 0.27 | -0.03 | -0.18 | 0.08 |
|  | non-supportive parenting behaviors | cognition | 0.21 | -0.02 | 0.19 | -0.003 | -0.087 | 0.09 |
|  | non-supportive parenting behaviors | language | 0.21 | 0.02 | 0.23 | 0.003 | -0.086 | 0.08 |
| Low Disadvantage (N=64) | **supportive parenting behaviors** | **cognition** | **-.49\*\*\*** | **.69\*\*\*** | **-0.08** | **-0.34\*\*\*\*** | **-0.54** | **-0.17** |
|  | **supportive parenting behaviors** | **language** | **-.49\*\*\*** | **.54\*\*** | **-0.31** | **-0.27\*\*** | **-0.53** | **-0.07** |
|  | **non-supportive parenting behaviors** | **cognition** | **.48\*\*\*** | **-.49\*** | **-0.18** | **-0.24\*** | **-0.42** | **-0.03** |
|  | **non-supportive parenting behaviors** | **language** | **.48\*\*\*** | **-.40\*** | **-0.4** | **-0.17\*** | **-0.38** | **-0.01** |

## DISCUSSION

PSD was nonlinearly associated with cognition and language scores at age 2 such that increasing PSD negatively predicted outcomes to a point, beyond which additional PSD was not associated with further decrements. Disadvantage moderated the relationships between parenting behaviors and CaL, such that parenting behaviors were only related to CaL for children with lower levels of disadvantage.

Supportive parenting behaviors at age 1 mediated the relationships of PSD to CaL. In contrast, brain metrics from birth did not mediate PSD-CaL associations. Critically, the mediating role of parenting in the association varied by level of disadvantage, which suggests that a basic level of resources may be necessary for parenting interventions to affect these key developmental outcomes. This finding has important implications for parenting interventions which are often applied universally, without consideration of psychosocial context. This critical finding should inform multi-step prevention and intervention efforts in early childhood for the greatest efficacy.

### PSD-related outcomes are detectable in toddlerhood

While *postnatal* socioeconomic status (SES) has reliably been associated with CaL, investigations of *prenatal* SD to CaL are sparse. Our finding that PSD was associated with lower language abilities is consistent with existing findings of poorer language scores in toddlers as a function of low *prenatal* SES.^8,47^ However, to our knowledge, this is the first report of *prenatal* disadvantage to cognition associations. We cannot rule out the contribution of postnatal SD to these associations, as pre- and postnatal SD were very stable in our sample and highly correlated (r=0.92). Nonetheless, our findings add to the growing literature documenting that the effects of SES begin very early in life,^48^ and highlight developmental consequences of PSD that are discernable as early as two years old. Importantly, our nonlinear findings also show that past a certain threshold of PSD, we do not detect significant associations of PSD with CaL. This nonlinearity may have important implications for understanding and addressing the effect of disadvantage on development, as it reveals that the effect of disadvantage varies by level. For example, interventions may fail to show CaL improvements if they do not first reduce disadvantage to a level where enhancing parenting behaviors could positively impact CaL in children, a consideration that is currently overlooked in most intervention programs.

### Supportive parenting behaviors mediate the relation of PSD to two-year outcomes; however, parenting has a stronger effect at lower levels of disadvantage

Supportive parenting mediated the relation of PSD to CaL. This is consistent with previous findings that similar parenting behaviors mediate the relationship of low SES to offspring outcomes, including reading, language, and working memory.^26–28^ As many of these mediations were observed in older samples, we extend the literature by demonstrating that parenting behaviors in early life can also mediate the relationship of *prenatal* disadvantage to toddler outcomes. While these findings may initially suggest that parenting behaviors in the first year of life may be a key target for intervention, we must also consider the finding that disadvantage moderated the relationship between parenting and neurocognitive outcomes.

Importantly, disadvantage moderated the relationship of parenting to CaL, such that greater supportive and fewer non-supportive parenting behaviors were related to higher CaL scores for children with *lower* levels of disadvantage. However, parenting was not related to CaL for children with higher levels of disadvantage. Further, our analyses revealed that both supportive and non-supportive parenting behaviors mediated the relation of PSD to CaL in the low disadvantaged group, *but no parenting behaviors mediated these relations in the high disadvantage group*. These findings are consistent with evidence that at varying levels of SES, different mechanistic pathways influence outcomes.^31^ These results suggest that while enhancing parenting practices (through socio-culturally informed^20,49,50^ and child specific ^51–53^ practices) may be helpful for children from advantaged background, it may be less effective when caregivers and children experience the highest levels of disadvantage. These effects need to be confirmed by causal study designs in the future, as they could underscore the dire need to provide families with a basic level of resources, as a minimum level of resources may be necessary for additional interventions (e.g., parenting) to improve child outcomes.

### Neonatal brain volumes do not mediate the relationship of PSD to cognitive and language abilities

While brain volumes were related to cognitive outcomes at year 2, we did not find evidence that birth brain metrics related to language scores or mediated PSD-CaL associations. Although neonatal structural brain volumes have been associated with age 2 language in prematurely-born children,^54^ the cortical development of preterm infants differs from that of term-born infants.^55^ Our absence of a strong association between neonatal structural volumes and language at age 2 in a full-term sample mirrors the null results of Spann et al^8^ in a smaller but similarly healthy sample in the same age range. As CaL are higher-order neurocognitive abilities, their neural correlates may not be developed enough at birth to be predictive of outcomes two years later. Relatedly, perhaps our ROIs (e.g., total cortical GM) were not specific enough to detect these associations, and analyses of smaller brain regions (i.e. perisylvian cortex^56^) would be more sensitive. Alternatively, a different imaging modality (e.g., resting state functional MRI) may be necessary to detect these associations.

### Strengths and Limitations

This study had several strengths, including a large sample of 202 mother-child dyads, a multidimensional conceptualization of *prenatal* disadvantage, investigations of nonlinear associations, and a longitudinal design to test mediation. However, there are also limitations. First, this study is correlational. As such, it is important to replicate these findings with future causal study designs to confirm whether supportive parenting and/or poverty reduction interventions would be effective for supporting cognitive and language development. Second PSD was highly correlated with disadvantage at year 1, so we could not investigate differential contributions of ongoing disadvantage. Third, parenting practices, and the perception of them by peers, assessors, and clinicians are sensitive to cultural beliefs and norms. As a large proportion of parenting research has been conducted with White participants^20^, the role and efficacy of parenting practices in different cultures has been less well characterized. As such, studies such as this one rely on valuations of parenting behaviors that may not be generalizable across cultures. Future research could greatly benefit from creating and using culturally inclusive parenting assessments. Further, all of the study’s PCI video coders were White, while 59% of participants were Black. While coders maintained high interrater reliability, future studies would benefit from assessment teams that match the demographic makeup of participants, which could foster cultural inclusivity in assessment practices.

## CONCLUSION

Collectively, these findings suggest that prenatal disadvantage exerts meaningful effects on key developmental outcomes in early life. Parenting behaviors constitute one mechanism through which this occurs; however, this mechanism may only be impactful at lower levels of disadvantage, suggesting that there may be a critical threshold beyond which mediating and moderating factors may become less effective. Thus, there is a dire need to address disadvantage as a primary prevention, bearing in mind that disadvantage encompasses multiple domains of social and material resources.

## List of Abbreviations

PSD: prenatal social disadvantage
CaL: cognition and Language
GM: grey matter
WM: white matter

