## Supplement for "Associations between parenting and cognitive and language abilities at age 2 depend on prenatal exposure to disadvantage"

This supplement includes:
Supplementary Text
Supplementary Tables eTable 1- eTable 2
Supplementary Figures eFigure 1
Supplementary References

### Supplemental Information

**Supplemental Text**


1. Prenatal Social Disadvantage Latent Variable
2. MRI Acquisition and Processing Pipeline
3. Detailed Study Exclusion Criteria
4. Parent-Child Interaction Task: Details, Coding Scheme, Coder and Composite Reliabilities
5. Analysis Covariates
6. Analysis Groupings for FDR correction
7. Interaction of Brain Volumes and Parenting


**Supplemental Tables**


**eTable 1**. The Association of Neonatal Brain Volumes with Cognition and Language Scores at Year 2

**eTable 2.** Relationships Between Prenatal Social Disadvantage and Year 2 Cognition and Language Are Not Mediated by Brain Volumes at Birth or by Non-Supportive Parenting Behaviors


**Supplemental Figures**


**eFigure 1**. Associations of Parenting Behaviors with Prenatal Social Disadvantage, Cognition, and Language

#### Supplemental Text

##### 1. Prenatal Social Disadvantage (PSD) Latent Variable.

*Maternal Observed Measures in Latent Variable, PSD.*[1] Home addresses were collected at delivery to calculate an Area Deprivation Index (ADI[2]) percentile. ADI indexes socioeconomic disadvantage at the neighborhood (e.g., census block) level using Census and American Community Survey data about education, housing quality, poverty, and employment. Health insurance status (private/public/no insurance) and highest education level were self-reported in the first trimester. Maternal nutrition status was obtained with the Healthy Eating Index.[3] An Income to Needs ratio[4] for each trimester was calculated based on self-reported income and household size standardized by the federal poverty threshold. For standardized estimates between the latent variable and observed components, see Luby et al.[1] Importantly, PSD does not include observed variables measuring maternal perceptions of prenatal psychosocial stress; and PSD and prenatal psychosocial stress (another latent variable created by the same group[1]) have dissociable effects on outcomes.[1,5,6]

##### 2. MRI Acquisition and Processing Pipeline.

As described in Triplett (2021),[5] T1- and T2-weighted and spin echo fieldmap data were acquired with the following sequence parameters, T1: repetition time (TR)=2400ms, echo time (TE)=2.22ms, voxel size=0.8×0.8×0.8 $mm^3$; T2: TR=3200/4500ms, TE=563ms, tissue T2=160ms, voxel size=0.8×0.8×0.8 $mm^3$, and spin echo: TR=8000ms, TE=66ms, voxel size=2×2×2 $mm^3$; 2 mm isotropic, multiband factor (MB)=1.

The T2-weighted images were first reviewed by a highly experienced imaging scientist (D.A.) and pediatric neurologist (C.D.S.) and evaluated based on image quality and estimated subject motion. Subjects determined to have severe motion during the scan (n=10) were not included in subsequent analyses.

The T2-weighted images were then preprocessed using the following standard steps: gradient and readout distortion correction using the Human Connectome Project preprocessing pipeline,[7] FSL axis reorientation to the MNI152 standard-space template,[8] image denoising using Advanced Normalization Tools for Brain and Image Analysis (ANTS) Registration Suite,[9] and co-registration using the Washington University School of Medicine Neuroimaging Laboratory (NIL)'s 4-dimensional floating point (4dfp)-based image analysis.[10] The resulting T2 images were then used as input for Melbourne Children's Regional Infant Brain atlas Surface (M-CRIB-S) segmentation and surface extraction toolkit, which automatically generated anatomical volume segmentations and reconstructed cortical surfaces.[11,12] The M-CRIB-S toolkit included N4 bias field correction and brain extraction, as well as automatic segmentation into white and gray matter, cerebellum, brainstem, and subcortical gray matter subdivisions corresponding to FreeSurfer-like labeling. Curvature-based spherical registration and mapping, alignment, and averaging were performed, allowing for spatial normalization within the cohort and to the M-CRIB atlas.

The segmentation volumes and the cortical surfaces were then projected on the T2 images (using Connectome Workbench[13] and ITK-SNAP[14] software packages) to qualitatively evaluate the concordance between segmentations and anatomic structures (including subcortical regions) and cortical surface reconstructions and anatomic gyral and sulcal morphometry. Segmentations and surfaces were rated independently by D.A. and a second, highly experienced rater (D.M.) for necessary edits as is standard with these analysis methods.[15–18] For a subset of subjects, segmentations were then manually edited (D.A. and D.M.) using the ITK-SNAP toolkit, and surfaces were regenerated using the M-CRIB-S toolkit. Edits were performed in all three planes to ensure accurate delineation of structures, primarily the supratentorial gray matter, white matter, and cerebrospinal fluid, also with minor edits of the subcortical structures and the cerebellum. Edited segmentations and surfaces were inspected iteratively, with additional minor edits, if necessary. Final segmentations and surfaces were reviewed and designated as complete by D.A. and C.D.S.

**3. Detailed Study Exclusion Criteria.**

The larger eLABE study recruited pregnant women without known pregnancy complications, infections known to cause congenital diseases, or positive drug screens (excluding tobacco and cannabis). The following conditions were used as exclusion criteria for neonates in the current study: preterm (born <37weeks GA), injury present on MRI birth scan, a NICU stay >7 days, specific NICU events (positive blood culture, antibiotics >3 days, intubated or chest tube, cooling or seizures, cardiac diseases, metabolic disorders), a birthweight of <2000g, maternal inflammation (e.g., oral or IV steroids).

**4. Parent Child Interaction Task.**

*Task Details.* The three tasks that parents completed with their child were: a Feeding task in which parent feeds child a snack, a Teaching task in which parent teaches child to find a ball under two cups, and a Free Play task in which parent and child freely play with a bin of toys. Caregivers were instructed to interact with their infant as they normally would in the home.

*Coding Scheme*: Each of the three parent-child interaction tasks were coded for the following parenting behaviors: Sensitivity, Positive Regard, Intrusiveness, Detachment, and Negative Regard. These were coded using a Parent-Child Interaction Rating Scale adapted from Brady-Smith et al.[19] All scales range from 1 to 7; higher scores indicate greater frequency and/or intensity of measure. Scores were averaged across all three tasks to maintain scaling.

- Sensitivity measures how interactions are child centered, well timed, responsive to cues, and appropriate to the needs, mood and capabilities of the child, and include actions such as acknowledging and matching affect, adjusting pace of play, and allowing the child to explore.
- Positive Regard measures expressions of positive affect, tone, warmth, enjoyment of, and praise for child, and involve taking into account consistency and range of behaviors.

- Intrusiveness measures the degree to which interactions are adult centered, overstimulating, and controlling, and include providing unwelcome suggestions, or denying the child the opportunity for autonomous play.
- Negative Regard measures expressions of anger and negativity toward the child, and includes behaviors ranging from disapproval and frustration to outward hostility and rejection.
- Detachment measures the lack of engagement, attention and awareness of the child, and includes behaviors ranging from lack of eye contact and flat affect to "checking out" and ignoring the child's cues for attention.

*Composite Details*. The Supportive Parenting Behaviors (SPB) composite averaged scores for sensitivity (e.g., noticing and responding to child cues) and positive regard (e.g., expressions of warmth, love, and praise). The Non-Supportive Parenting Behaviors (NSPB) composite averaged scores for intrusiveness (e.g., directive, controlling, or adult-centered interactions), detachment (e.g., lack of attention to and engagement with the child), and negative regard (e.g., expressions of anger, disapproval, or rejection of the child).

*Coder and Composite Reliabilities.* Reliability was established by coders reaching a "gold standard" with the master coder of 92% exact or within 1 before coding interactions independently, and 20% of tapes were coded for ongoing reliability. Intraclass correlation coefficients (ICCs) computed at the conclusion of the study all indicated good reliability (Average ICC > .70)[20]. Average ICCs are as follows: Sensitivity (ICC = .874), intrusiveness (ICC = .855), positive regard (ICC = .846), negative regard ( ICC= .780), detachment (ICC = .872)

**5. Analysis Covariates.**

The following variables were related to Bayley-III outcomes of interest, thus were included as covariates in all analyses.

- gestational age [cognition: r=.23, p=.001; language: r=.19, p=.007]
- age at assessment [cognition: r=.13, p=.033; language: r=.14, p =.06]
- sex [cognition: r=.10, p=.18; language: r=.16, p=.02]

**6. Analysis Groupings for FDR Corrections.**

The families for FDR corrections are as follows:

- Does prenatal social disadvantage (PSD) predict outcomes at year 2?
  - 3 tests; 1 correction family
    - (a) PSD → cognition

        - (a) PSD → language
        - (a) PSD → motor
- Do PSD-associated neonatal brain volumes predict outcomes at year 2?
    - 12 tests; 2 correction families
        - (a) cortical GM → cognition scores
        - (a) subcortical GM → cognition scores
        - (a) cerebral WM → cognition scores
        - (a) cerebellar volumes → cognition scores
        - (a) total BV → cognition scores
        - (a) mean gyrification index → cognition scores

        -----------------------------

        - (b) cortical GM → language scores
        - (b) subcortical GM → language scores
        - (b) cerebral WM → language scores
        - (b) cerebellar volumes → language scores
        - (b) total BV → language scores
        - (b) mean gyrification index → language scores
- Does PSD predict parenting behaviors at year 1?
    - 2 tests; 1 correction family
        - (a) PSD → SPB
        - (a) PSD → NSPB
- Do parenting behaviors predict outcomes at year 2?
    - 4 tests; 2 correction families
    - (a) SPB → cognition scores
    - (a) NSPB → cognition scores

    ------------------

    - (b) SPB → language scores
    - (b) NSPB → language scores

**7. Interactions between brain volumes and parenting behaviors.**

We also tested whether brain metrics and parenting interacted to predict outcomes by regressing brain metrics, parenting, the interaction of brain metrics and parenting, and covariates onto outcomes. There were no significant interactions between disadvantage-associated brain volumes and parenting behaviors (supportive or non-supportive) in predicting year 2 outcomes.

**eTable 1. The association of neonatal brain volumes with cognition and language scores at year 2.**

| | Cognition | | | Language | | |
|---|---|---|---|---|---|---|
| **Brain Volumes at Birth** | **β** | ***p*** | ***q*** | **β** | ***p*** | ***q*** |
| Total cortical gray matter | 0.13 | 0.68 | 0.68 | 0.07 | 0.40 | 0.40 |
| Total cerebral white matter | **0.23** | **< 0.01** | **0.02** | 0.13 | 0.11 | 0.22 |
| Total cerebellum | 0.13 | 0.08 | 0.13 | 0.21 | 0.18 | 0.21 |
| Subcortical gray matter | **0.19** | **0.01** | **0.03** | 0.16 | 0.04 | 0.22 |
| Total brain volume | **0.18** | **0.01** | **0.03** | 0.11 | 0.15 | 0.22 |
| Mean GI | 0.08 | 0.340 | 0.41 | 0.11 | 0.17 | 0.22 |

**PSD-associated brain volumes at birth are related to cognition but not language outcomes at year 2.**
Standardized estimates (ß), *p*-values, and *q*-values (FDR-corrected *p*-values) are indicated. *P*-values were corrected within each outcome (e.g., 2 groups of 6 tests). Covariates were sex, age, and GA. **Bolded effects are significant after correction.**

**eTable 2. Relationships between prenatal disadvantage and year 2 cognition and language are not mediated by (A) brain volumes at birth or (B) non-supporting parenting behaviors**

#### A. Mediations by Neonatal Brain Volumes

| *Mediating Variable* | *Outcome* | *Covariates* | *path A* | | *path B* | *direct effect* | *indirect effect* | *95% CI Indirect Effect* | |
|---|---|---|---|---|---|---|---|---|---|
| | | | *Δ Adj $R^2$* | *EDF* | *β* | *β* | *β* | *Lower* | *Upper* |
| total WM | cognition | sex, age, GA | 15.3%**** | 1.40 | 0.08 | -0.04 | -0.03 | -0.080 | 0.020 |
| subcortical GM | cognition | sex, age, GA | 12.2%**** | 1.40 | 0.06 | -0.05 | -0.02 | -0.061 | 0.030 |
| total brain volume | cognition | sex, age, GA | 13.6%**** | 1.40 | 0.04 | -0.06 | -0.01 | -0.080 | 0.030 |

#### B. Mediations by Parenting Behaviors

| *Mediating Variable* | *Outcome* | *Covariates* | *path A* | | *path B* | | *direct effect* | *indirect effect* | *95% CI Indirect Effect* | |
|---|---|---|---|---|---|---|---|---|---|---|
| | | | *Δ Adj $R^2$* | *EDF* | *Δ Adj $R^2$* | *EDF* | *β* | *β* | *Lower* | *Upper* |
| Non-supportive parenting behaviors | cognition | sex, age, GA | 29.62%**** | 1.00 | -4.5% | 1.00 | -0.160 | -0.089 | -0.240 | 0.020 |
| Non-supportive parenting behaviors | language | sex, age, GA | 29.62%**** | 1.00 | -0.4% | 1.55 | -0.041 | -0.075 | -0.193 | 0.020 |

**Relations between prenatal disadvantage and year 2 cognition and language are not mediated by (A) brain volumes at birth, or (B) non-supportive parenting behaviors.**

**Change in adjusted $R^2$ ( Δ Adj. $R^2$), expected degrees of freedom (EDF) and GAM *p* values for paths modeled with GAMS (RLRT *p* < .05). Standardized effect estimates (β ) for paths modeled linearly, direct and indirect effects**. Change in adjusted R2 is the difference between the adjusted $R^2$-values of a model with and without the predictor of interest, where positive values indicate that the predictor of interest improves the model fit. Path A describes the effect of the independent variable (prenatal disadvantage) to the mediator variable, controlling for sex, age, and gestational age at birth (GA). Path B describes the unique effect of the mediator variable on the dependent variable, controlling for sex, age, and GA. Direct effect indicates the unique effect of independent variable (prenatal disadvantage) on the dependent variable, while controlling for the mediating variable, sex, age, and GA. The indirect effect is the mediation effect.[21]
**No mediations are significant.** *p* < .05, ***p* < .01, ****p* < .001, *****p* < .0001

**eFigure 1. Associations of Parenting Behaviors with Prenatal Social Disadvantage, Cognition, and Language**

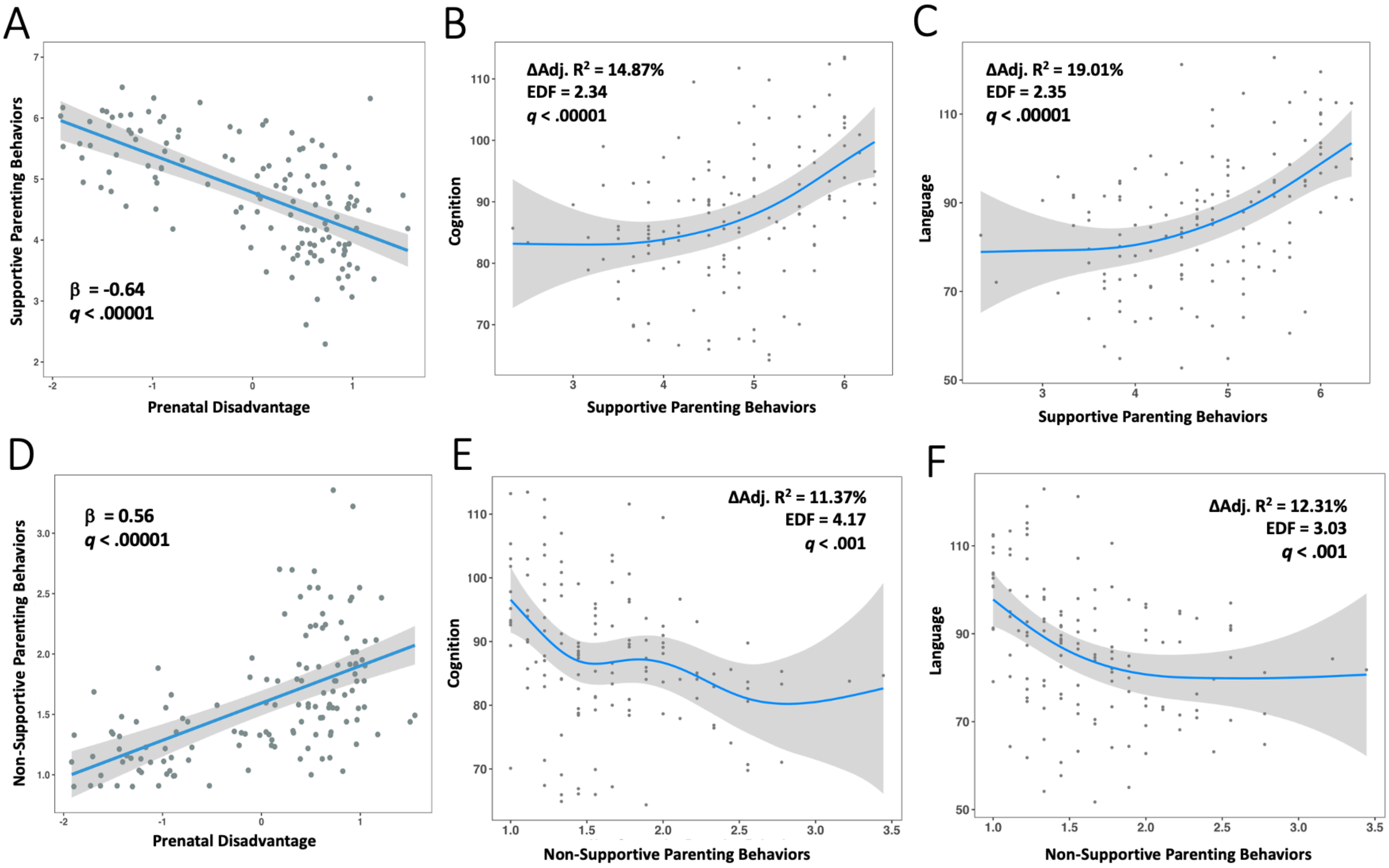


Supportive parenting behaviors at year 1 are (A) linearly associated with prenatal social disadvantage (RLRT p's > .05) and (B, C) nonlinearly (RLRT *p's* < .05) associated with outcomes cognition and language scores at year 2, whereby the relationship of supporting parenting to each outcome was stronger with higher supportive parenting scores. Non-Supportive parenting behaviors at year 1 are (D) linearly associated with prenatal social disadvantage (RLRT p's > .05) and (E, F) nonlinearly (RLRT *p's* < .05) associated with outcomes cognition and language at year 2, whereby such that the relation of non-supportive parenting to cognition and language was strongest at lower levels of non-supportive parenting. All figures illustrate the partial effect of independent variable (e.g., controlling for covariates sex, age, and GA) on dependent variable. EDF = expected degree of freedom for the shape of the smooth. Change in Adjusted $R^2$($\Delta$ Adj. $R^2$) = the difference between the adjusted $R^2$-values of a model with and without the predictor of interest, where positive values indicate that the predictor of interest improves the model fit.
